# Sex and stress interactions in fear synchrony of mouse dyads

**DOI:** 10.1101/2024.06.09.598132

**Authors:** Wataru Ito, Alexei Morozov

## Abstract

Socially coordinated threat responses support the survival of animal groups. Given their distinct social roles, males and females must differ in such coordination. Here, we report such differences during the synchronization of auditory-conditioned freezing in mouse dyads. To study the interaction of emotional states with social cues underlying synchronization, we modulated emotional states with prior stress or modified the social cues by pairing unfamiliar or opposite-sex mice. In same-sex dyads, males exhibited more robust synchrony than females. Stress disrupted male synchrony in a prefrontal cortex-dependent manner but enhanced it in females. Unfamiliarity moderately reduced synchrony in males but not in females. In dyads with opposite-sex partners, fear synchrony was resilient to both stress and unfamiliarity. Decomposing the synchronization process in the same-sex dyads revealed sex-specific behavioral strategies correlated with synchrony magnitude: following partners’ state transitions in males and retroacting synchrony-breaking actions in females. Those were altered by stress and unfamiliarity. The opposite-sex dyads exhibited no synchrony-correlated strategy. These findings reveal sex-specific adaptations of socio-emotional integration defining coordinated behavior and suggest that sex-recognition circuits confer resilience to stress and unfamiliarity in opposite-sex dyads.

## Introduction

Behavioral coordination in social organisms, especially when responding to threats, is necessary for group survival, requiring the rapid integration of cues representing potential threats and social cues from conspecifics. Studies show that in animals and also in humans, social cues affect emotional states, which, in turn, alter the perception of these cues (Taylor 1981, Sandi and Haller 2015, Muroy, Long et al. 2016), suggesting an intertwining of the brain circuits responsible for processing affective and social information.

Several forms of socially modulated threat responses have been characterized in rodents (Morozov and Ito 2019). They include enhanced fear learning after exposure to stressed peers (Knapska, Mikosz et al. 2010), observational fear learning (Chen, Panksepp and Lahvis 2009, Jeon, Kim et al. 2010), fear conditioning by proxy (Bruchey, Jones and Monfils 2010, Jones, Riha et al. 2014), and fear buffering by non-fearful conspecifics (Kikusui, Winslow and Mori 2006, Kiyokawa, Takeuchi and Mori 2007, Guzman, Tronson et al. 2009). These forms, however, reflect delayed and unidirectional social effects, which are assessed mostly in a non-social context by testing single subjects individually or several days after the social interactions. In contrast, socially coordinated threat responses entail real-time, bidirectional cue exchange among animals responding to immediate threats simultaneously. This aspect of social behavior remains overlooked in current studies.

We developed a paradigm for studying coordinated threat responses in mice (Ito, Palmer and Morozov 2023), where auditory fear-conditioned mouse dyads exposed to a continuous tone (120 s) as a conditioned stimulus (CS) show synchronized freezing patterns. The affective CS drives freezing in each mouse, but synchronization depends on social cues from the partner. This setup allows examination of how affective and social elements interact and shape behavioral synchronization. We observed more robust synchronization in male mice than females (Ito, Palmer and Morozov 2023), prompting further investigation into sex differences in fear synchrony.

Sex differences in behavioral coordination, as shown in various human studies (Weitz 1976, Lafrance and Ickes 1981, Tronick and Cohn 1989, Fujiwara, Kimura and Daibo 2019), likely reflect the distinct roles of males and females in handling social group threats (Van Vugt, De Cremer and Janssen 2007). Given the female’s primary role in nurturing and protecting progeny, Taylor and colleagues proposed that, unlike males, females evolved to “selectively affiliate in response to stress, which maximizes the likelihood that multiple group members will protect them and their offsprings” (Taylor, Klein et al. 2000). Given the well-documented sex differences in social and emotional processing (Taylor, Klein et al. 2000, Reyes, Valentino and Van Bockstaele 2008, Bangasser, Reyes et al. 2013, Bangasser and Cuarenta 2021), we hypothesized that the interplay between social and affective brain systems varies between sexes.

We examined this by altering affective states and social information in the fear synchrony paradigm. Dyads were given mild stress to change affective states and assembled with familiar or unfamiliar partners and same or opposite-sex partners to vary social cues.

The modulations resulted in, first, sex-specific effects on fear synchrony in same-sex dyads, implying sex differences in the interplay between the social and emotional systems. Second, synchrony in opposite-sex dyads was resilient to stress and unfamiliarity, suggesting a unique and potentially adaptive mechanism that modulates the affective system, differentiating the responses to social cues depending on the partners’ sex.

## Results

### The social and non-social components of fear synchrony

We recently reported that mice in dyads synchronize conditioned freezing (Ito, Palmer and Morozov 2023); however, whether the synchronization relies exclusively on social cues has not been examined. Suppose two mice showed similar temporal freezing dynamics due to intrinsic freezing properties or any reason; our synchrony metric interprets these similar dynamics as a high degree of synchrony, even if it has nothing to do with exchanging social cues. Therefore, we investigated the non-social synchrony component through the following two approaches.

First, we compared synchrony between partners tested in the same chamber (PAIR) and in separate chambers isolated from each other without exchanging social cues (SINGLE). We fear-conditioned the mice and ran the two tests the following day. To mitigate the impact of repeated testing, we separated the two tests by a 5-hour interval and counterbalanced the testing sequences between “PAIR➜SINGLE” and “SINGLE➜PAIR” (**Fig 1a**). The synchrony was significantly above 0 in PAIR (male: p<0.0001, female: p<0.001) but not SINGLE configuration in both sexes (**Fig1b**), suggesting a negligible non-social component.

**Fig 1.**
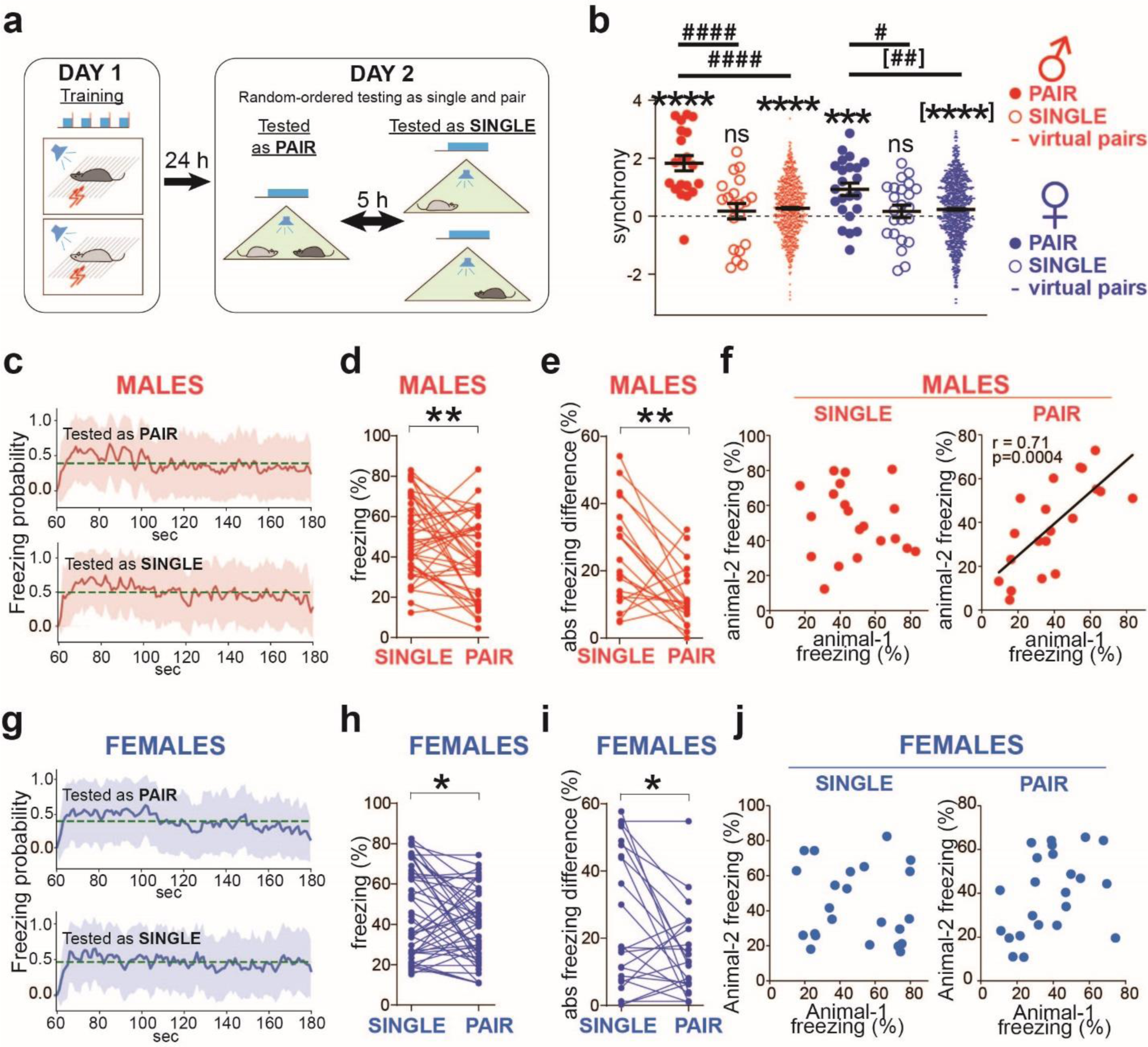
Male dyads coordinate freezing more strongly than females. **a** Scheme for testing the social modulation of freezing. Fear conditioning on Day 1, followed by counterbalanced sequential fear testing in either the ‘PAIR➜SINGLE’ or ‘SINGLE➜PAIR’ orders on Day 2. **b** Significant synchrony when tested together (PAIR) but not in isolation (SINGLE). However, the virtual pairs reveal a small non-social synchrony component. Horizontal bars indicate means±SEM. One-sample t-test comparing to 0: *** p<0.001, **** p<0.0001. Wilcoxon Signed Rank Test comparing to 0: [****] p<0.0001. Two sample t-test: ^#^ p<0.05, ^####^ p<0.0001. Mann-Whitney test: [^##^] p<0.01. Male dyads n=20, female dyads n=23, virtual male dyads n=760, virtual female dyads n=1012. **c, g** Averaged freezing probability during the CS period, computed using a sliding window of 1s. Dashed lines show the probability means. Shadowed areas represent SD (male n=40, female n=46). **d, h** Freezing levels of individual animals tested alone (SINGLE) or with a partner (PAIR). n=40 (males), 46 (females). Wilcoxon matched pairs test: * p<0.05, ** p<0.01. **e, i** Absolute freezing differences between partners tested individually (SINGLE) and together (PAIR). Male (n=20) and female (n=23) dyads. Wilcoxon matched pairs test: * p<0.05, ** p<0.01. **f, j** Freezing levels of male partners correlated when tested together (PAIR) but not separately (SINGLE), and the correlations in females were not significant in both cases. The black lines show linear regression. The correlation coefficients (r) and p-values are shown in the case of statistically significant correlations.

Nevertheless, it remained a concern that individually tested mice exhibited different freezing patterns from when tested with partners because of the isolation. To exclude such isolation effect, we computed synchrony in the virtual dyads – all possible combinations between mice tested as PAIR excluding real experimental pairs (20 male and 23 female real dyads yielded 760 male and 1012 female virtual dyads). It revealed small non-social synchrony components (**Fig 1b**, virtual pairs, males: 0.27±0.03, females: 0.23±0.03, significantly positive due to overpowering by high N) and similar to the synchrony in the SINGLE configuration (males 0.17±0.27, females: 0.17±0.21). Both types of non-social synchrony were significantly smaller than the synchrony in the PAIR configuration (**Fig 1b**, males: p<0.0001, PAIR compared to both SINGLE configuration and virtual pairs; females: p=0.02, PAIR compared to SINGLE, p=0.0011, PAIR compared to virtual pairs). Meanwhile, computing the dynamics of the averaged freezing probabilities revealed a similar temporal pattern in both testing configurations: a rapid initial rise in about 5 seconds after the CS onset, followed by a slow decline regardless of the partner’s presence or sex (**Fig 1c, g**), presumably the cause of the identified small non-social synchrony components.

Finally, we examined whether mice synchronize to a non-social object – the Sphero Mini Robot Ball (Sphero, Hong Kong) (Solie, Contestabile et al. 2022), programmed to mimic precisely the freezing patterns from previously tested mice. The ball moves continuously during the pre-CS period and then stops and occasionally moves during CS, as the tested mice did (**Supplementary Fig 1**, left). The mean synchrony with the ball was positive but not significantly different from 0 in both sexes (**Supplementary Fig 1**, clean ball, males: 0.29±0.28; females: 0.42±0.22). To endow the ball with a social characteristic, we coated it with urine from a mouse of the opposite sex relative to the subject. Both sexes showed significant synchrony with the urine-pained ball (**Supplementary Fig 1**, urinated ball, males: 0.79±0.30, p=0.024; females: 1.20±0.37, p=0.015).

Those three results indicate that the observed synchronized freezing is driven primarily by social cues in both sexes. Nevertheless, we updated our synchrony quantification by subtracting the non-social component computed from the virtual dyads in every experimental group. In addition, male dyads exhibited higher synchrony than females (p=0.0095) (**Fig 1b**), which replicated our earlier findings (Ito, Palmer and Morozov 2023).

### Equalization of fear response is more pronounced in males than in females

We further investigated the influences of the conspecific presence on fear levels and possible sex effects. In line with previous studies (Kiyokawa, Takeuchi and Mori 2007, Fuzzo, Matsumoto et al. 2015), mice of both sexes demonstrated fear buffering by decreasing freezing in the partner’s presence, with this effect being more significant in males (p=0.007) than females (p=0.029) (**Fig 1d, h**). Additionally, the absolute difference between partners’ freezing levels was smaller in both sexes when tested as PAIR than when tested as SINGLE. Notably, this equalization was more significant in males (p=0.008) than in females (p=0.032) (**Fig 1e, i**). Furthermore, partners’ freezing levels did not correlate during testing as SINGLE (**Fig 1f, j**, left SINGLE) but correlated strongly in males tested as PAIR (p=0.0004) and not in females (**Fig 1f, j,** right PAIR). These data indicate that social cues equalize fear levels more strongly in male than female dyads.

### Sex-specific behavioral strategies underlying high synchrony

To identify behavioral processes underlying sex differences in fear synchrony, we first examined freezing properties in both sexes, computing three parameters: the averages of freezing levels, freezing bout duration, and freezing bout counts. Although we did not find sex differences by comparing each parameter directly (**Fig 2a,c,e**), testing the correlations with fear synchrony revealed parameter-specific sex differences. The averages of freezing levels and duration of freezing bouts correlated with synchrony in females but not males (**Fig 2b,d**). Contrarily, the number of freezing bouts correlated with synchrony in males but not females (**Fig 2f**). The sex-specific correlations suggested that males and females synchronized using distinct behavioral strategies.

**Fig 2.**
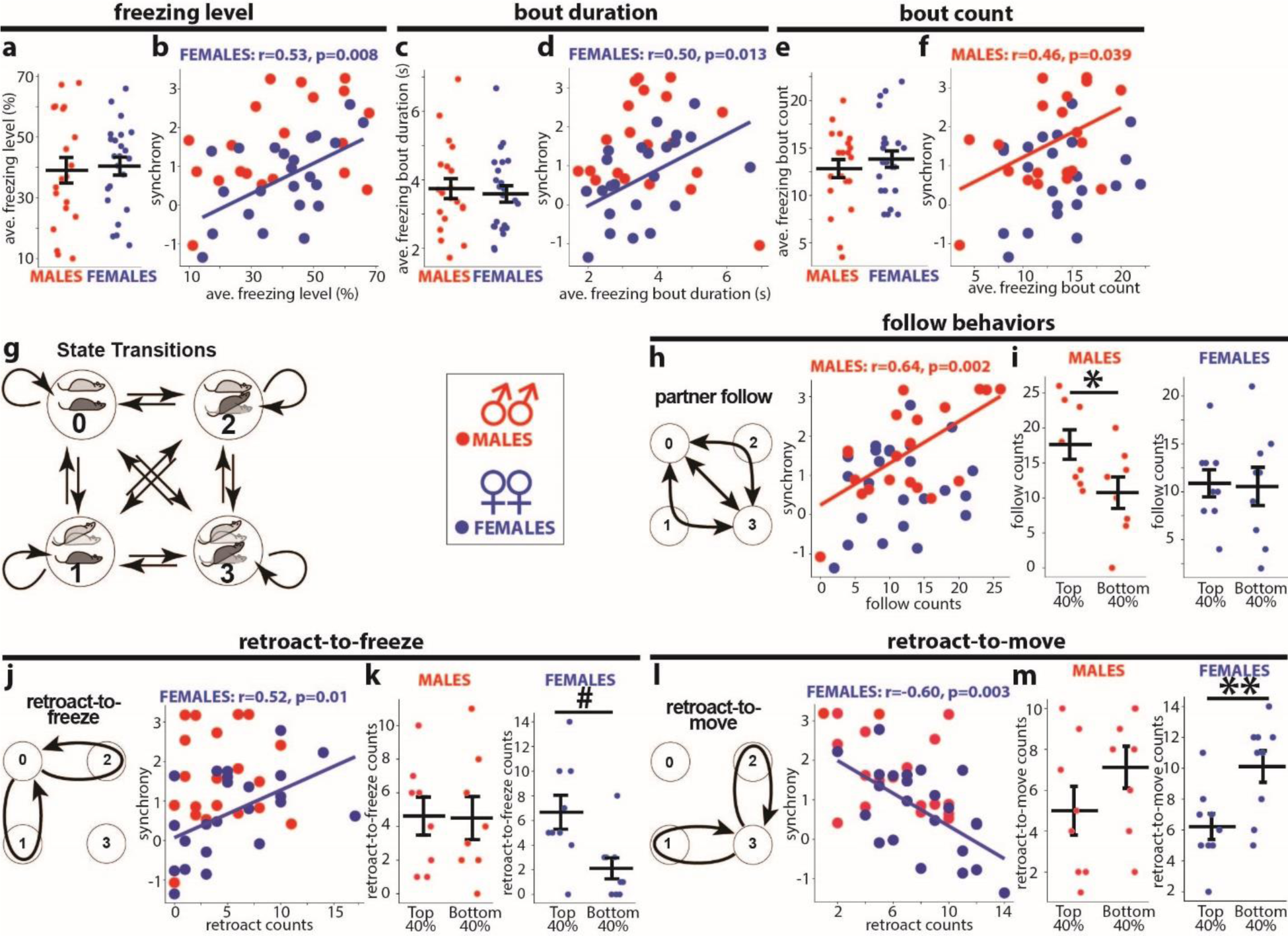
Sex-specific behavioral correlates of synchrony. **a-f** Freezing parameters (**a,c,e**) and their scatter plots with synchrony (**b, d, f**): dyad’s average freezing (**a**), freezing bout duration (**c**), and freezing bout counts (**e**). **g** Four behavioral states of a dyad (state 0,1,2,3): horizontal and angled mouse symbols represent freezing and moving mice, respectively. Arrows depict all possible transitions among the four states. **h** “Follow” behaviors. Left: Scheme of the “follow” behaviors as transitions between states 0 and 3 directly or via intermediate states 1 and 2. Right: Scatter plots between synchrony and the “follow” counts. **i** Comparisons of the “follow” counts between the top and bottom 40% dyads ranked by synchrony. **j, l** “Retroact-to-freeze” (**j**) and “retroact-to-move” (**l**) schemes and scatter plots as in **h**. **k,m** The same comparisons as in **i**. Dyads numbers: males: n=20, females: n=23. The Pearson correlation coefficients (r), significance values (p), and trend lines are shown for significant correlations. Horizontal bars on diagrams indicate means±SEM. *p<0.05, **p<0.01 two-sample t-test, #p<0.05, Mann-Whitney test.

Next, we decomposed the synchronization process into intricate behavioral states and subsequent transitions. We used a Markov chain model, describing the dynamic of two dyad members, animal A and B, as transitions among four unique states, states 0, 1, 2, and 3 (**Fig 2g**). In state 0, animals A and B both freeze. In states 1 and 2, one animal freezes while the other moves. In state 3, both animals move. We designated states 0 and 3 as ‘congruent,’ symbolizing the same actions, while states 1 and 2 were labeled ‘incongruent,’ representing contrasting actions. State transitions count a total of 16, including four self-looping transitions. We obtained complete time series data of the state transitions from each dyad based on manual video analyses of freezing bouts.

We then focused on two distinct behavioral processes predicted to influence fear synchrony: the “follow” and the “retroact” behaviors. The “follow” behaviors can be formalized as two-step transitions between the congruent states 3 and 0 via the non-congruent states 1 or 2 or directly (**Fig 2h left**). We define one as “follow-to-freeze” when one mouse starts freezing from state 3, and the partner follows. Conversely, if one mouse begins to move from state 0, and the other follows, we define it as “follow-to-move.” In all dyads analyzed, the counts of “follow-to-freeze” and “follow-to-move” were equal or differed by 1; therefore, we used their sums to represent the “follow” behaviors.

Meanwhile, the “retroact” behaviors can be formalized as two-step loops that start in the congruent states 0 or 3, pass via states 1 or 2, but return to the starting state (**Fig 2j,l left**). The “retroact” behaviors include two types: One is “retroact-to-freeze,” in which one mouse breaks the state 0 by moving and then reverses itself to freezing (retroacts), returning the dyad to state 0. Another is “retroact-to-move,” in which one mouse breaks state 3 by freezing and reverses itself to moving (retroacts), returning the dyad to state 3. The “retroact-to-freeze” favors synchrony, bringing animals to simultaneous freezing in state 0. In contrast, the “retroact-to-move” opposes synchrony by decreasing the chance of state 0.

These decomposed behavioral processes showed no sex differences in the occurrence or counts (data not shown). However, they exhibited sex-specific correlations with synchrony. The “follow behaviors” counts positively correlated with synchrony in males but not females (**Fig 2h right**). Accordingly, the top 40% of male dyads ranked by synchrony had higher “follow” counts than the bottom 40% (**Fig 2i left**), but females did not show such differences (**Fig 2i right**). Contrarily, female synchrony correlated positively with the “retroact-to-freeze” counts (**Fig 2j right**) and negatively with the “retroact-to-move” (**Fig 2l right**), but males exhibited no such correlations. Accordingly, the top 40% ranked female dyads had higher counts in “retroact-to-freeze” and lower counts in “retroact-to-move” than those within the bottom 40% (**Fig 2k, m right**). By contrast, males did not exhibit such differences. The negative correlation of “retroact-to-move” counts is consistent with its unfavorable effects on synchrony.

These results indicate that the distinct behavioral processes, the “follow” and “retroact” behaviors, are the sex-specific determinants of the level of fear synchrony in male and female dyads, respectively, and represent sex-specific behavioral strategies for achieving high synchrony. Back to the freezing properties, these strategies can partially explain their sex-specific correlations with synchrony (**Fig 2b,d,f**). The higher “follow” counts require more freezing bouts, explaining their correlation to synchrony in male dyads (**Fig 2f**). Likewise, the longer freezing bouts allow higher “retroact-to-freeze” counts, explaining the correlation between freezing bout durations and synchrony in females (**Fig 2d**).

### Mild stress diminishes fear synchrony in male dyads but augments it in females

To induce a negative affective state, we subjected both dyad members to 5-minute restraint stress one hour before the synchrony test (Lolait, Stewart et al. 2007, Newson, Pope et al. 2013) (**Fig 3a**). For synchrony, the two-way ANOVA revealed a significant stress*sex interaction (F_(1,75)_ = 19.9, p<0.0001). The stressed males’ synchrony was lower than in non-stressed males (p=0.0008) and did not differ from zero (**Fig 3b left**). Unexpectedly, the stressed females’ synchrony was higher than in non-stressed females (p=0.0037) and well above zero (p<0.0001) (**Fig 3b right**). For freezing levels, the two-way ANOVA revealed a significant stress*sex interaction (F_(1,154)_ = 11.3, p=0.001). Freezing levels did not differ between stressed and non-stressed males but were higher in stressed than non-stressed females (p<0.0001) (**Fig 3c**). Moreover, the partners’ freezing levels no longer correlated with each other in stressed males, as in female dyads, regardless of stress (**Supplementary Fig 2, Fig 1j**).

**Fig 3.**
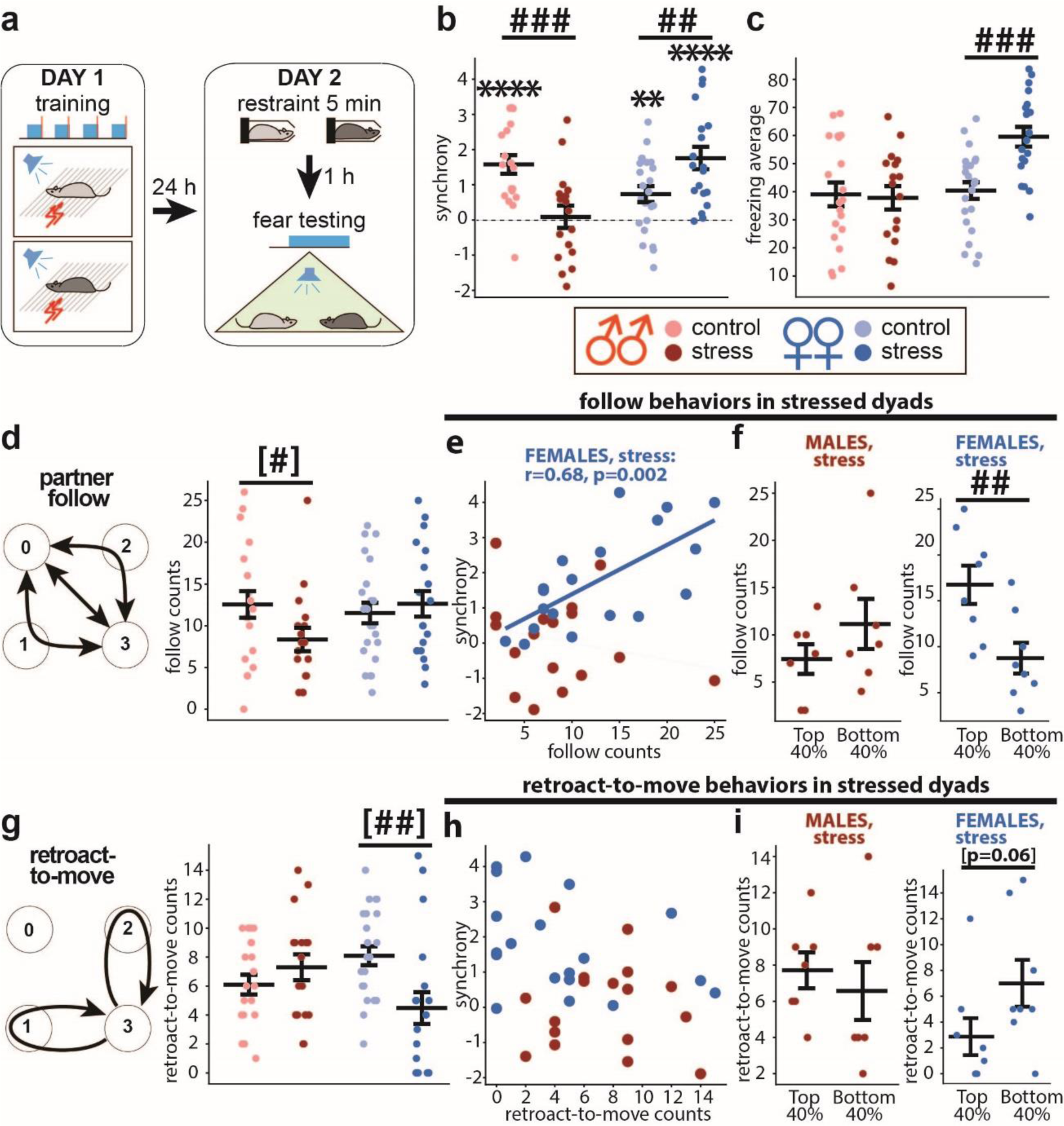
Sex-dimorphic effect of stress on fear synchrony and the underlying behaviors. **a** Scheme for testing stress effect on fear synchrony. Fear conditioning training on Day 1, and five min restraint followed by fear testing after 1h on Day 2. **b, c** Summary diagrams for synchrony (**b**) and freezing (**c**). Colors representing sex (red: male dyads, blue: female dyads) and treatments (light symbols: control, males: n=20, females: n=23, dark symbols: stress, males: n=17, females: n=19). The color scheme is the same in the following panels. **d “**Follow” behavior scheme and the counts in control and stressed dyads. **e** Scatter plot between synchrony and “follow” counts in stressed dyads. **f** Comparisons of “follow” counts between the top and bottom 40% of stressed dyads ranked by synchrony. **g-i** “Retroact-to-move” scheme and the count data are shown as in **d-f.** The Pearson correlation coefficients (r) and significance values (p) are shown only for significant correlations. Horizontal bars on diagrams indicate means±SEM. One-sample t-test comparing to 0: ** p<0.01, **** p<0.0001. Two-sample t-test: ^##^p<0.01, ^###^p<0.001. Mann-Whitney test: ^[#]^p<0.05, ^[##]^p<0.01.

The occurrence of “follow behaviors” was decreased by stress in males but not females (**Fig 3d**). Consistently, the “follow” counts no longer correlated to synchrony in stressed males but opposingly correlated in stressed females (**Fig 3e**). It appeared that stress switched the sex-specific correlation patterns. Accordingly, the top 40% of stressed female dyads ranked by synchrony exhibited more “follow” counts than the bottom 40% (**Fig 3f right**). For the “retroact-to-freeze,” stress did not affect its counts in either sex but abolished the correlation to synchrony in females (**Supplementary Fig 3a,b**). For the “retroact-to-move” behavior, the stressed females decreased their counts (**Fig 3g right**) and lost the synchrony correlation to the counts (**Fig 3h**). Accordingly, there were no differences in the counts between the top and bottom 40% dyads ranked by synchrony (**Fig 3i right**). The lower ‘retroact-to-move’ counts and higher synchrony in stressed females align with their negative correlation in control females ( **Fig 2l**). In summary, stress decoupled the “follow” behaviors from high synchrony in male dyads but, conversely, linked them in female dyads.

### Inactivation of the dorsomedial prefrontal cortex protects males’ synchrony from stress

Recent studies have shown that adverse experiences in mice disrupt social behaviors by activating the dorsomedial prefrontal cortex (dmPFC) (Xu, Liu et al. 2019, Huang, Zucca et al. 2020). Based on these findings, we hypothesized that dmPFC mediated the stress-induced disruption of fear synchrony in male dyads. Since the dmPFC inactivation by muscimol did not affect fear synchrony in non-stressed males (Ito, Palmer and Morozov 2023), we tested the same dmPFC inactivation on stressed male dyads (**Fig 4a, b**). Muscimol increased synchrony (p=0.008) (**Fig 4c left**) without changing the freezing levels (**Fig 4c right**) and also increased the “follow” behaviors count (**Fig 4d**). The “follow” counts did not correlate to synchrony in the stress+vehicle group, the same as in **Fig 3e,** but correlated in the muscimol group (**Fig 4e**). The top 40% of dyads ranked by synchrony exhibited more “follow” counts than the bottom 40% in the muscimol but not the vehicle group (**Fig 4f**). In summary, dmPFC muscimol inactivation in stressed males restored synchrony and the link between the “follow behaviors” and high synchrony.

**Fig 4.**
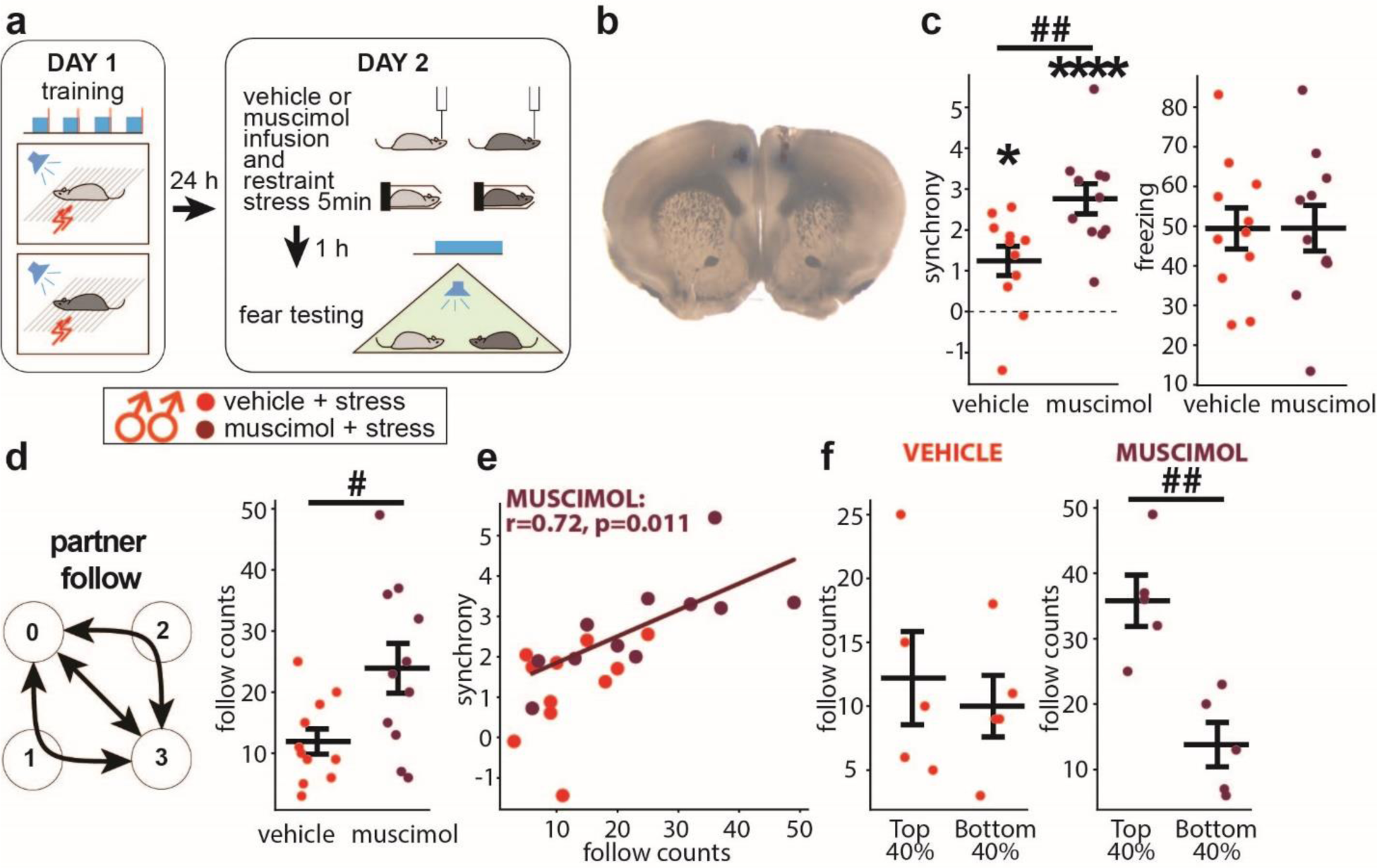
dmPFC inactivation increases synchrony in stressed males. **a** Scheme for testing dmPFC inactivation on fear synchrony in stressed male dyads (n=11 (vehicle, light red), 11 (muscimole, dark red)). **b** An example brain slice with the muscimol injection site in dmPFC identified by Chicago Sky Blue injection. **c** Summary diagrams for synchrony (**left**) and freezing (**right**). **d “**Follow” behavior scheme and comparison of the counts. **e** Scatter plot between synchrony and “follow” counts. **f “**Follow” counts comparisons between the top and bottom 40% of dyads ranked by synchrony. The Pearson correlation coefficients (r) and significance values (p) are shown only for significant correlations. Horizontal bars on diagrams indicate means±SEM. One-sample t-test comparing to 0: *p<0.01, ****p<0.0001. Two-sample t-test: ^#^p<0.05 ^##^p<0.01.

### Unfamiliarity decreases fear synchrony in males and has sex-specific effects on freezing

As components of social context, the partner’s familiarity and sex interact with emotional transmission between partners. Prior studies have shown that fear is modulated more effectively among familiar rodents (Jeon, Kim et al. 2010, Kiyokawa, Honda et al. 2014, Pisansky, Hanson et al. 2017). Moreover, familiar male mice exhibit a more robust fear transmission than females (Pisansky, Hanson et al. 2017). Accordingly, we examined if the same rules applyed to fear synchrony. As predicted, unfamiliar male dyads displayed lower synchrony than familiar ones (p=0.04, one-tailed t-test), but no differences were found in females (**Fig 5a**).

**Fig 5.**
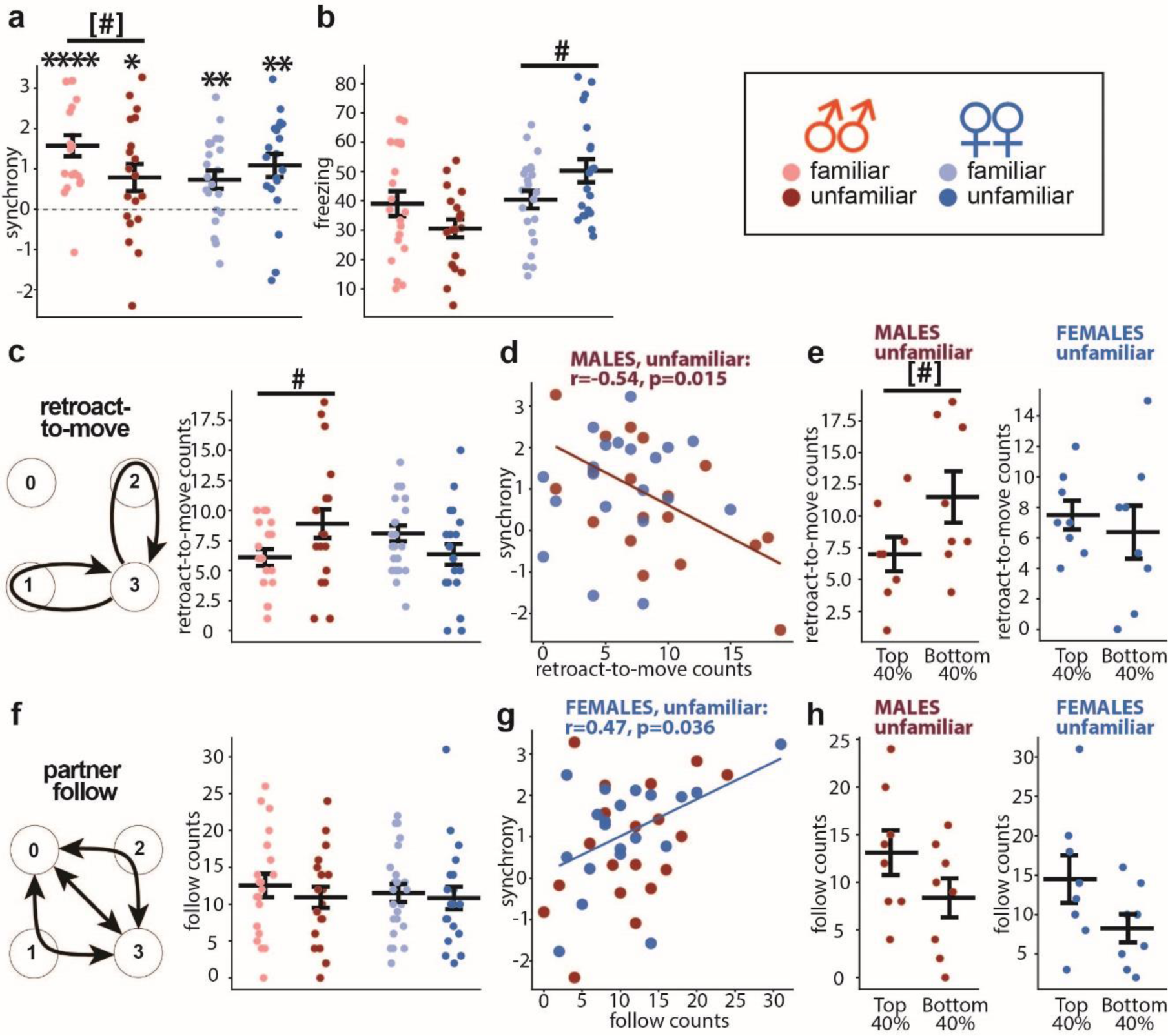
Unfamiliarity decreases fear synchrony in males. **a** Synchrony in males (red) and females (blue), familiar (light symbols, male dyads, n=20, female dyads, n=23) and unfamiliar (dark symbols, male dyads, n=19, female dyads, n=20). **b** Freezing levels. **c “**Retroact-to-move” scheme (**left**) and summury of the counts (**right**). **d** Scatter plot between synchrony and “retroact-to-move” counts in unfamiliar dyads. **e “**Retroact-to-move” counts comparisons between the top and bottom 40% of unfamiliar dyads ranked by synchrony. **f “**Follow” behavior scheme and comparison of the counts. **g** Scatter plot between synchrony and “follow” counts. **h “**Follow” counts comparisons between the top and bottom 40% of dyads ranked by synchrony. The Pearson correlation coefficients (r) and significance value (p) are shown only for a significant correlation. Horizontal bars on diagrams indicate means±SEM. One sample t-test compared to 0: *p<0.05, **p<0.01, ****p<0.0001, two-sample t-test: ^#^p<0.05 (two-sided), ^[#]^p<0.05 (one-sided).

Unfamiliarity increased freezing levels in female dyads (p=0.049) (**Fig 5b**) and abolished the correlation between freezing and synchrony (data not shown), as seen in familiar females (**Fig 2b**). Meanwhile, unfamiliarity did not change freezing levels in males (**Fig 5b**) but abolished freezing equalization between partners as the loss of correlation between their freezing levels (data not shown). Females never showed such a correlation, regardless of the partners’ familiarity.

Moreover, decomposing behaviors revealed that, in males, unfamiliarity abolished the correlation of synchrony to the “follow” counts without changing the mean of the counts (**Fig 5f,g**). At the same time, unfamiliarity increased the “retroact-to-move” counts (p=0.046) (**Fig 5c**) and enabled their negative correlation to synchrony (**Fig 5d**), resulting in fewer “retroact-to-move” counts in the top 40% than the bottom 40% dyads ranked by synchrony (p=0.043, one-tailed t-test) (**Fig 5e**).

In contrast, unfamiliarity had the opposite effects in females, eliminating the correlation between synchrony and the “retroact-to-move” counts (**Fig 5d**) and causing a moderate correlation of synchrony to the “follow” counts (r=0.47, p=0.036) (**Fig 5g**); however, it did not cause statistically significant differences in the “follow” counts between the top 40% and the bottom 40% dyads ranked by synchrony (**Fig 5h**). Both changes occurred without changing the mean of each count (**Fig 5c,f**).

### No impact from stress or unfamiliarity in opposite-sex dyads

Finally, we examined fear synchrony with opposite-sex partners and tested the effects of stress and unfamiliarity. The “familiar” opposite-sex dyads comprised mice housed together for 5-7 days before training and testing. For the “stress” group, the familiar dyads underwent a 5 min immobilization one hour before testing, as outlined in **Fig 2a**. In contrast, the “unfamiliar” group consisted of mice housed with same-sex cagemates throughout the training and tested with an unfamiliar partner of the opposite sex.

Fear synchrony was highly significant across all three groups, with no discernable differences among groups (**Fig 6a**). There were no differences in freezing levels, freezing bout counts or durations between the groups (data not shown). The freezing levels showed the correlation between partners only in the familiar group without stress (**Fig 6c**), indicating that stress and unfamiliarity attenuated fear equalization, similar to their effects in male dyads.

**Fig 6.**
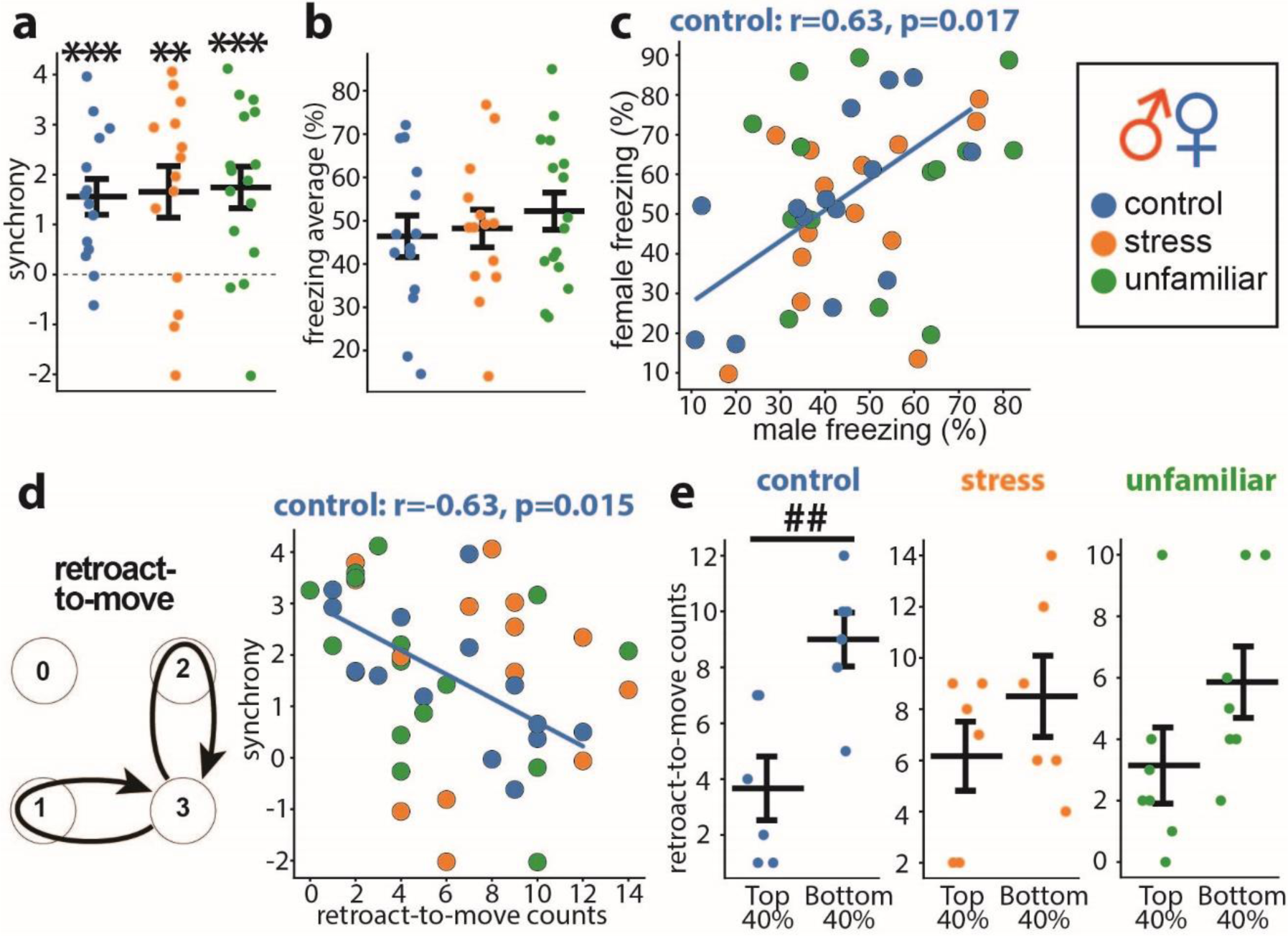
No stress and familiarity effects on male-female dyads. **a** Synchrony in male-female dyads of familiar control (blue, n=14), stressed (orange, n=14), and unfamiliar animals (green, n=16). The color coding of groups is the same in all panels. **b** Average freezing. **c** Scatter plots of the female freezing as the function of the male freezing show correlation only between familiar unstressed partners (blue: controls). **d** “Retroact-to-move” scheme and scatter plots between synchrony and “retroact-to-move” counts. **e “**Retroact-to-move” counts comparisons between the top and bottom 40% of dyads ranked by synchrony in each group. The Pearson correlation coefficients (r) and significance value (p) are shown on the scatter plot for a significant correlation. Horizontal bars on diagrams indicate means±SEM. One-sample t-test: **p<0.01, ***p<0.001. Two-sample t-test: ^##^p<0.01.

The only behavioral process that correlated with synchrony was “retroact-to-move.” The negative correlation was observed only in familiar dyads without stress (**Fig 6d**). Accordingly, the top 40% of such dyads ranked by synchrony exhibited fewer counts than the bottom 40% (p=0.005) (**Fig 6e**). The lack of a correlation of synchrony with the “follow “ and “retroact-to-freeze” counts indicates that the opposite-sex dyads use both behavioral processes equally to achieve high synchrony.

## Discussion

This study demonstrates that mice synchronize auditory conditioned freezing in male, female and opposite-sex dyads. We identified sex-specific behavioral strategies for high synchrony in male and female same-sex dyads, which differentiated the effects from prior mild stress and the partner’s unfamiliarity. Intriguingly, the opposite-sex dyads were resilient to both stress and unfamiliarity, lacking preferred strategies.

We examined two forms of socially coordinated freezing: 1) coordinated temporal patterns of freezing bouts, or “fear synchrony” (Ito, Palmer and Morozov 2023), and 2) coordinated levels of freezing, or “freezing equalization.” The effects of the emotional state and social cues were tested by giving mild stress and pairing unfamiliar or opposite-sex partners. Four key findings emerged. 1) In familiar same-sex non-stressed dyads, males coordinated freezing more strongly than females. Meanwhile, the opposite-sex dyads coordinated similarly to the levels of the male dyads. 2) Male and female dyads used different behavioral strategies to achieve high synchrony: the “follow” and “retroact-to-freeze” behaviors, respectively, whereas the opposite-sex dyads exhibited no preferred strategy. 3) Mild stress abolished the freezing synchrony of male dyads in a dmPFC-dependent manner but instead facilitated it in females and had no effect on the opposite-sex dyads. 4) Pairing with unfamiliar partners attenuated synchrony in male dyads, not females and opposite-sex dyads.

Before investigating sex differences, we tested how much non-social factors could impact the synchrony readout, driven by a concern that a temporal similarity of slow freezing dynamics contributes to spurious synchronization independent of social cues. Eliminating social cues by testing animals in separate chambers, computing synchrony in virtual dyads, or testing synchronization with a non-social object revealed a minor non-social synchrony component. Accordingly, the synchrony computation was updated by subtracting the non-social component determined using the virtual dyads method.

As sex differences in control same-sex dyads, we first replicated the lower fear synchrony in females than males (Ito, Palmer and Morozov 2023). Second, we also found weaker coordinated levels of freezing in females, revealed by the following metrics: 1) less significant fear buffering (**Fig 1d, h**) (Kiyokawa, Takeuchi and Mori 2007, Fuzzo, Matsumoto et al. 2015), 2) less significant freezing equalization (**Fig 1e, i**), and 3) non-significant correlation between partners’ freezing levels, which was significant in males (**Fig 1f, j**).

To delineate the behavioral causes for the sex differences, we sought the behavioral metrics that determine the synchrony. First, we investigated the freezing properties—freezing levels, freezing bout durations, and the bout counts. Despite these metrics did not differ between sexes, they exhibited sex-specific correlations to synchrony. The freezing bout counts positively correlated to synchrony only in males (**Fig 2f**), whereas the freezing levels and the bout durations - only in females (**Fig 2bd**), suggesting that sex-specific behavioral processes drive synchrony.

Next, we decomposed the behavioral dynamics of dyads using the state transition analysis based on a Markov chain model. Then we identified two types of behavior processes relevant to synchrony: 1) the “follow” behavior, in which partners copy each others’ transitions between freezing and non-freezing, switching the dyad between the two congruent states 0 and 3, and 2) the “retroact” behavior, in which one mouse breaks a congruent state but reverses the action after not followed by the partner, returning the dyad to the initial congruent state, either freezing (retroact-to-freeze) or moving (retroact-to-move). While the mean counts of these behaviors did not differ between the sexes, they exhibited sex-specific correlations to synchrony. In males, the counts of “follow” behaviors correlated positively (**Fig 2h**), whereas, in females, the “retroact-to-freeze” and “retroact-to-move” showed positive and negative correlations, respectively (**Fig 2j,l**).

Fear synchrony arises from an innate motivation to freeze in unison with a partner, which is evident when dyad members’ freezing bouts overlap more frequently than by chance, regardless of sex. However, the identified sex-specific behavioral correlations suggest that each sex prefers a different process to achieve high synchrony. These two processes vary in assigning the roles of “synchrony breaker” and “synchrony fixer” between partners: during the “follow” behavior, each member takes one of these roles, whereas during the “retroact” behavior, a single mouse takes both. When dyads transition from congruent to incongruent states, the “synchrony breaker” changes its freezing state, reflecting emotional alteration, and the remaining member observes the breaker’s action. The subsequent choice of “synchrony fixer” could depend on how the two members differentially perceive each other. In the “follow” process, preferred by males, the “synchrony breakers” ignore their partner’s state, leaving the fixing role to the partner to bring the dyad into a congruent state. By contrast, in the “retroact” process, preferred by females, the “synchrony breakers” attend to the partner and reverse their actions to bring the dyad into a congruent state.

We reported that the projection from the ventral hippocampus (vHPC) to the basal amygdala (BA) mediates fear synchrony (Ito, Palmer and Morozov 2023), presumably conveying the freezing state of the partners. Opposingly, the BA has a robust reciprocal projection to vHPC, possibly conveying the freezing state of the self. When the “synchrony breakers” change their freezing state, the BA➜vHPC projection could modulate social neuronal ensembles in the vHPC representing the partner’s freezing states, potentially underlying the sex difference in the effects of emotional switches on social perception in the “synchrony breakers.”

Mild stress effects on fear synchrony contrasted between sexes. Stress wiped out fear synchrony in males but facilitated it in females, suggesting the presence of sex-specific regulatory mechanisms. Accordingly, we found that dmPFC-inactivation protected male dyads from stress-induced loss of synchrony (**Fig 4**). Interstingly, the dmPFC is not required for fear synchrony in control males (Ito, Palmer and Morozov 2023), indicating that stress recruits dmPFC to interfere with male synchrony, which is consistent with the reported dmPFC-dependent disruptions of social behaviors by stress (Xu, Liu et al. 2019, Huang, Zucca et al. 2020). The decomposed synchrony process revealed that the stressed male dyads decreased the “follow” behavior counts and lost their correlation to synchrony.

Meanwhile, mild stress raised synchrony in females to the levels seen in the control males. Interestingly, the stressed females lost the synchrony-correlations of the “retroact-to-freeze” count but gained the correlations of the “follow” count, seen in the control male dyads, indicating that stress switched females to the males’ strategy for achieving high synchrony. In addition, stressed females decreased the “retroact-to-move” count, consistent with the high synchrony due to a higher chance of both mice freezing.

The exact molecular and circuit mechanisms of stress-enhanced female synchrony remain unknown. However, several stress-responsive molecules (Bangasser and Cuarenta 2021) could play a role. For instance, oxytocin, released in response to stress (Bangasser and Cuarenta 2021), promotes prosocial behaviors in females, possibly due to estrogen-enhancing oxytocin functions (McCarthy 1995). Another candidate is the CRF-1 receptor in the locus coeruleus (LC), which shows sex-dependent regulation: it desensitizes in males but remains active in females (Bangasser, Reyes et al. 2013), sustaining LC activity and elevated arousal. In addition, the larger size of the female’s LC and its longer dendrites can underly the sustained female’s arousal and attention after stress (Bangasser, Eck et al. 2018). A recent study using mice’s operant social stress paradigm revealed that males and females use different stress-coping behavioral strategies. While males exhibit social avoidance, females pursue social interaction (Navarrete, Schneider et al. 2024), which can be explained by stress-induced social arousal and is consistent with our finding of elevated fear synchrony in stressed females. Moreover, the lack of stress-induced synchrony loss, as seen in males, can result from the protective effects of estrogen in females, which dampens the dmPFC response to stress (Wei, Yuen et al. 2014, Velli, Iordanidou et al. 2022).

Unfamiliarity moderately reduced fear synchrony and diminished freezing equalization in males (**Fig 5a**). These findings agree with previous reports of reduced social modulation by unfamiliar male conspecifics in several fear paradigms, including observational fear learning, fear conditioning by proxy, and fear buffering (Jeon, Kim et al. 2010, Kiyokawa, Honda et al. 2014, Pisansky, Hanson et al. 2017, Agee, Jones and Monfils 2019). Conversely, unchanged synchrony in unfamiliar females aligns with the report on females’ familiarity-independence of observational fear learning (Pisansky, Hanson et al. 2017). The sex difference in the unfamiliarity effects can arise from different social perceptions of same-sex conspecifics. A male mouse naturally living in the harem perceives an unfamiliar male as a threat rather than a partner, which likely prevents defensive coordination. In contrast, the minimal intrasexual competition in females (Pandolfi, Scaia and Fernandez 2021) likely allows defensive coordination between strangers.

It was intriguing to test the effects of opposite-sex partners on fear synchrony, a unique non-reproductive social behavior. The control opposite-sex dyads exhibited high synchrony, the same as the male same-sex dyads, indicating that social cues from a male partner elicit in females more robust coordination than cues from another female in same-sex dyads. Moreover, synchrony in the opposite-sex dyads was resilient to stress and unfamiliarity, indicating that social cues from a female partner counteract the adverse effects of stress and unfamiliarity in males. Decomposing synchrony processes revealed no synchrony-correlations in the “follow” or “retroact” behaviors, except the negative correlation to “retroact-to-move” in the control dyads, suggesting that the opposite-sex dyads achieve high synchrony using both behavioral strategies.

The primary social cues in mice are the chemosensory signals transmitted via the vomeronasal pathway, activating sex-encoding neurons in the medial amygdala, hypothalamus, and prefrontal cortex (Li and Dulac 2018, Kingsbury, Huang et al. 2020). These signals further reach the circuits underlying coordinated defensive behaviors, such as the anterior hypothalamus and medial preoptic area, which can drive affiliation (Fukumitsu, Kaneko et al. 2022, Jabarin, Dagash et al. 2023), as well as the ventral hippocampus (vHPC) and basal amygdala (BA), which mediate fear synchrony (Ito, Palmer and Morozov 2023). In the case of opposite-sex dyads, the sex-recognition circuits appear to modulate the socio-emotional integrator underlying fear synchrony in both partners. In females, the modulation makes the integrator more responsive to the male partner, whereas in males, it confers resilience to stress. Investigating the roles of these circuits in fear synchrony can unravel how the brain differentially integrates socio-emotional cues depending on the sex of the information source.

In this study, we extended the scope of the fear synchrony paradigm, a real-time social modulation of emotional states represented by freezing bouts, to the sexual domains. As a non-reproductive social behavior, the model allowed us to investigate same-sex and opposite-sex dyads. It revealed sex-specific synchronization strategies in same-sex dyads and the effects of the sex-recognition circuit. The decomposed synchronization dynamics will help tackle the neuronal substrates of that complex behavior in future studies.

## Materials and methods

All experiments were performed according to a Virginia Tech IACUC-approved protocol.

### Animals

Breeding trios of one C57BL/6N male and two 129SvEv females produced 129SvEv/C57BL/6N F1 hybrid male and female mice weaned at p21 and housed four littermates per cage of the same sex as described (Ito, Erisir and Morozov 2015). Subjects underwent tests at p75-p90. The same-sex familiar dyads were formed from the littermates 7 days before training and placed in cages with fresh bedding, which was not changed until completion of the testing. For the same-sex unfamiliar dyads, the mice were housed in pairs of littermates for 7 days before testing but paired randomly with unfamiliar non-littermates from different cages during testing. The familiar opposite-sex dyads comprised mice housed together for 5-7 days before training and testing from the pool of the virgin animals, initially housed in groups of 4 same-sex littermates per cage. The unfamiliar opposite-sex dyads consisted of mice housed permanently in groups of same-sex littermates. We provide the entire list of animal groups in this study (**Supplementary Table 1**).

### Fear conditioning

Mice in each dyad were trained independently in two separate conditioning chambers (Med Associates, St. Albans, VT) and then tested together as dyads in a single chamber as described (Ito, Palmer and Morozov 2023). For training, each animal received four pairings of the conditioned stimulus (CS: a 30 s, 8 kHz, 80 dB tone) and unconditioned stimulus (US: a 0.5 mA 0.5 s electrical shock co-terminated with CS) given in variable intervals (60-180 s). Cued fear was tested one or two days later in a new context on a dyad (PAIR) or a single animal (SINGLE) per chamber. The animals spent the first 1 min without CS and then 2 min with CS. Videos were recorded at four frames per second, exported as AVI files with MJPEG compression using the Freezeframe system (Actimetrics, Wilmette, IL), and then converted to the mp4 format using a Python script.

### Immobilization stress

Mice were placed inside the 50 ml Falcon tubes with an 8 mm opening at the bottom near the nose of subjects to allow breathing for 5 min situated outside of the home cage. Then, the animals were immediately returned to the home cage.

### Quantification of freezing, freezing Synchrony and partners’ freezing correlations

Freezing was defined as the absence of all observable movement of the skeleton and vibrissae, except for those related to respiration (Fanselow 1980), during at least four consecutive video frames (1s). Briefly, annotators, unaware of the treatment of the animals, manually identified the first and last video frames of each freezing bout and recorded them using a Python script. From the recorded annotation data, another Python script computed the freezing duration for each animal and then freezing overlap, chance overlap, and freezing synchrony for each dyad. Freezing synchrony was defined as the standardized difference between the observed and chance freezing overlaps (Ito, Palmer and Morozov 2023) minus the non-social synchrony component, computed as the mean synchrony in the virtual dyads that included all possible pairs between mice of the given group except the real pairs.

Before testing the correlation between partners’ freezing in the same-sex dyads, we used reshuffling to avoid a spurious correlation due to unbalanced freezing levels between the correlated groups. We first sorted the dyads by the mean freezing levels, second, identified the higher and lower freezer animals in each dyad, and third, performed alternated assignment of the high or low freezers as animal 1 or animal 2 moving sequentially from the dyad with the highest mean freezing to the one with the lowest mean freezing. The resulting animal 1 and animal 2 groups were balanced by freezing means and numbers of high and low freezers. For the opposite-sex dyads, we computed correlations between male and female freezings.

### Quantification of copying and self-correction behaviors

From the annotation data in the previous section, the behavioral states of each dyad were sampled with every video frame (4 frames per second), giving 480 samples during 2 min of the CS period. The “follow” behavior counts were computed as the total number of transitions between state 0 and state 3, directly or via states 1 or 2, in either direction. The “retroact-to-freeze” and “retroact-to-move” counts were computed as the total number of the two-step state 0➜1/2➜0 and state 3➜1/2➜3 loops, respectively.

### Muscimol inactivation of dmPFC

The inactivation by local muscimol infusion was performed as described (Ito, Palmer and Morozov 2023). Briefly, 10-14 days before fear conditioning, mice received bilateral implantation of the guide cannula (P1 Technologies, Roanoke, VA) at 1.5 mm anterior, ±0.5 mm lateral from bregma, and −0.8 mm ventral from the brain surface. Dummy cannulas were placed in the guide cannulas to prevent clogging. Animals were handled for 2-3 minutes daily, including removal and reattachment of dummy cannulas, during the 7 days before testing. One day before fear conditioning, mice were habituated to infusion using the vehicle: (in mM) 150 NaCl, 10 D-glucose, 10 HEPES, 2.5 KCl, 1 MgCl_2_, pH 7.35. The infusion cannula extended 1 mm over the guide cannula. The infusion volume was 155 nL per site, and the infusion rate was 75 nL/min. Muscimol (1.17 mM) or vehicle was infused 1 hour before fear testing. After the final test, mice were anesthetized with 2.5% Avertin and received an intracranial infusion of Chicago Sky Blue, 0.2 % (Sigma-Aldrich), followed by transcardial perfusion with 4% paraformaldehyde. Visible light microscopy verified the infusion sites.

### Data analysis

Statistical analyses were performed using GraphPad Prism 10 (GraphPad Software, La Jolla, CA). Normality was tested using the Shapiro–Wilk test. Datasets with normal distribution were compared using the one-sample or two-sample t-test. The datasets with non-normal distribution were compared using the Wilcoxon signed-rank test and Mann– Whitney test. All the tests were two-sided except when mentioned otherwise. The two-tailed p-value was calculated for the Spearman or Pearson correlation analyses. The effects were deemed significant with p < 0.05.

## Data and code availability

All primary data, including video files, are available from authors upon reasonable request. In addition, codes for data analysis and statistics will be provided with example data as part of the replication package after publication.

## Acknowledgments

The study was supported by an NIH grant R01MH120290 to AM.

---

**Supplementary Fig 1.**
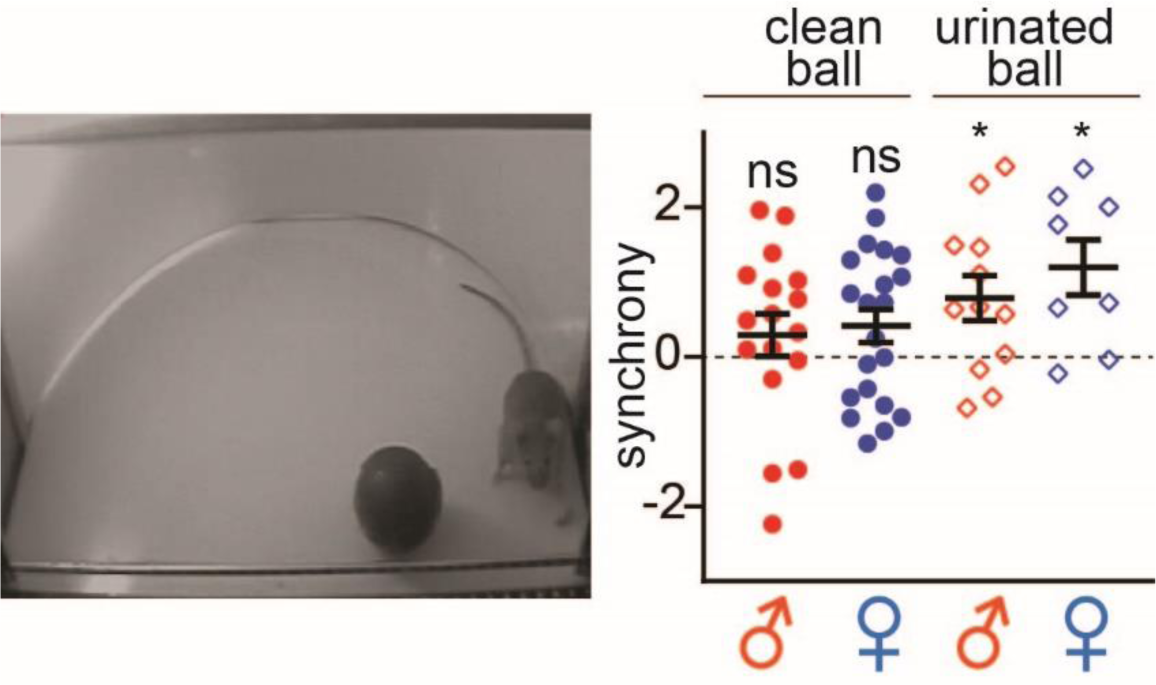
Mice do not synchronize with non-social ball, but urine-painting on the ball enables synchrony. **Left** Synchrony test with non-social ball. **Right** Summary of synchrony for each group (clean ball: n=17 (male), 21 (female), urinated ball: n=12 (male), 8 (female)). One-sample t-test comparing to 0: * p<0.05.

**Supplementary Fig 2.**
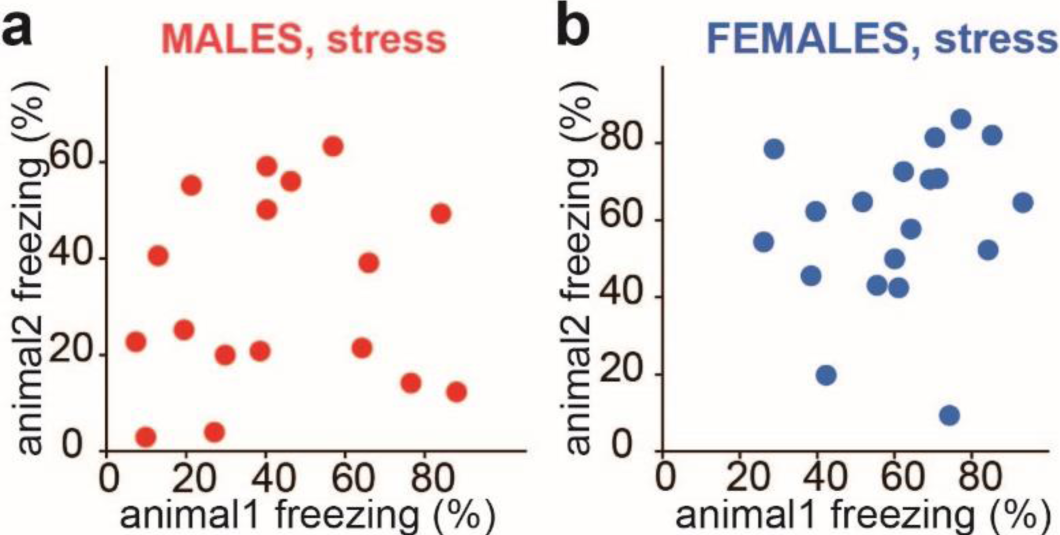
Freezing levels of stressed partners do not correlate. Scatter plots of partners’ freezing levels in male (**a**) (n=17) and female (**b**) (n=19) dyads.

**Supplementary Fig 3.**
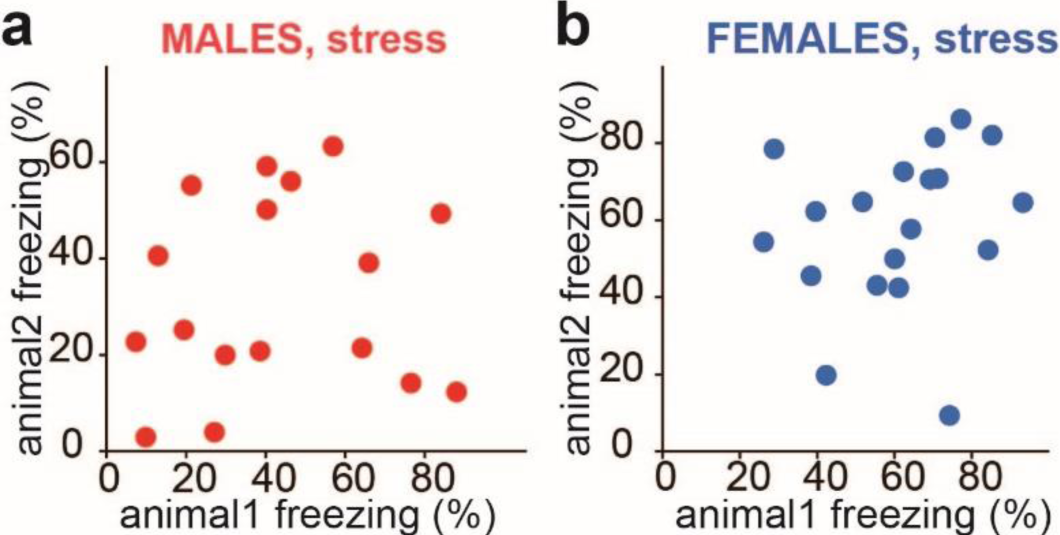
Stress effects on retroact-to-freeze. **a** “Retroact-to-freeze” scheme (**left**) and counts in control and stressed dyads (**right**). **b** Scatter plot between synchrony and “retroact-to-freeze” counts in stressed dyads. **c** Comparisons of **”**Retroact-to-freeze” counts between the top and bottom 40% of stressed dyads ranked by synchrony. Colors representing sex (red: male dyads, blue: female dyads) and treatments (light symbols: control, males: n=20, females: n=23, dark symbols: stress, males: n=17, females: n=19). Horizontal bars on diagrams indicate means±SEM.

**Supplementary Table 1.**
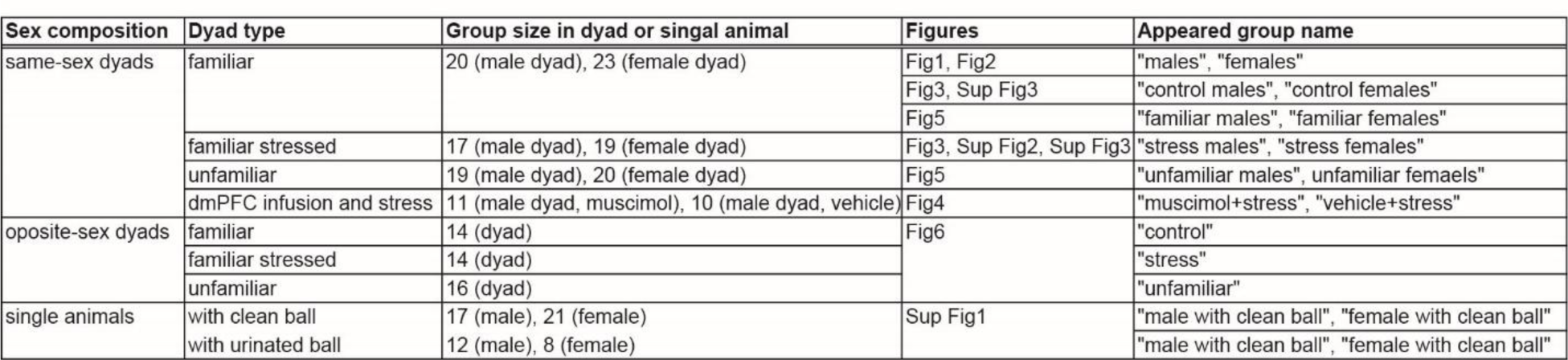
The list of dyad groups in the study.

